# Distinct contributions of hippocampal pathways in learning regularities and exceptions revealed by functional footprints

**DOI:** 10.1101/2025.09.09.674568

**Authors:** Melisa Gumus, Michael L. Mack

## Abstract

Fundamental aspects of learning are theorized to be supported by hippocampal pathways: monosynaptic pathway (MSP) extracts regularities whereas trisynaptic pathway (TSP) rapidly encodes exceptional items. Yet, the empirical evidence for the dynamic involvement of MSP and TSP in learning remains elusive. We leveraged diffusion-weighted imaging to estimate the end points of MSP- and TSP-related white matter structures (i.e., footprints) within hippocampal subfields and entorhinal cortex. We then measured the activation of pathway-specific footprints with functional magnetic resonance imaging while participants learned novel concepts defined by regularities and exceptions. The functional footprint method revealed links between MSP-related footprint activation and regularity encoding early in learning, and TSP-related footprint activation and exception encoding late in learning. These findings provide novel evidence that learning concept regularities and exceptions is distinctly supported by hippocampal pathways. Pathway footprint approach provides insights into the functional dynamics of the human hippocampus, translating theoretical and computational work into empirically testable questions in humans.

**Significance Statement:** Theories suggest that learning involves two main functions of the hippocampus: integrating commonly encountered information and distinctly encoding exceptional items. Yet, the theorized contributions of hippocampal pathways are yet to be empirically validated. We provide the first evidence of this in humans by introducing a multimodal technique (pathway footprints) that targets learning-related activations at the endpoints of hippocampal pathways. The more a pathway is engaged for a given learning experience, the more the pathway endpoints will be activated. This innovation reveals the dynamic involvement of hippocampal pathways in learning: one unites common information early in learning while another encodes exceptional items late in learning. The pathway footprint method offers precise estimates of neural signatures of cognition as refined by anatomy.

## 1. Introduction

One fundamental paradox of memory is the transformation of singular moments into a stable scaffold of knowledge. While memories are anchored in time and space (Tulving, 1972), modern evidence suggests that the hippocampus, the canonical seat of episodic memory, also builds structured knowledge, simultaneously preserving and generalizing from distinct individual experiences (Brunec et al., 2020; Eichenbaum, 2017; Sherman et al., 2023). Such flexible memory arises from the hippocampus and its anatomically distinct subfields. Indeed, innovative approaches in animal models and humans have demonstrated key evidence of flexible memory in hippocampal function and representations including pattern separation of similar events (Bakker et al., 2008; Leutgeb et al., 2007), integration of experiences that share elements (Larkin et al., 2014; Schlichting et al., 2014), abstraction of regularities across related experiences (Schapiro et al., 2012), and goal-dependent coding of the same sensory information (Mack et al., 2016). To achieve these complex behaviours, influential computational work (Schapiro et al., 2017; Sučević & Schapiro, 2023) delineates the complementary roles of two central hippocampal pathways: the monosynaptic pathway (MSP), which directly connects entorhinal cortex (ERC) to Cornu Ammonis 1 (CA1), extracts and generalizes regularities over many repetitions, while the trisynaptic pathway (TSP), which loops in dentate gyrus (DG) and CA2/3, rapidly binds together elements to learn distinct episodes (Fig. 1a) (Norman & O’Reilly, 2003; Schapiro et al., 2017). A key untested prediction from this theoretical framework is that the relative engagement of the two hippocampal pathways impacts what is learned from a given experience. Specifically, whereas MSP supports the extraction of regularities shared across experiences, TSP contributes to distinct encoding of the current experience. Here, we evaluate this theoretically driven hypothesis in humans with a novel method for measuring the functional engagement of hippocampal pathways during a flexible concept learning task.

**Fig. 1:**
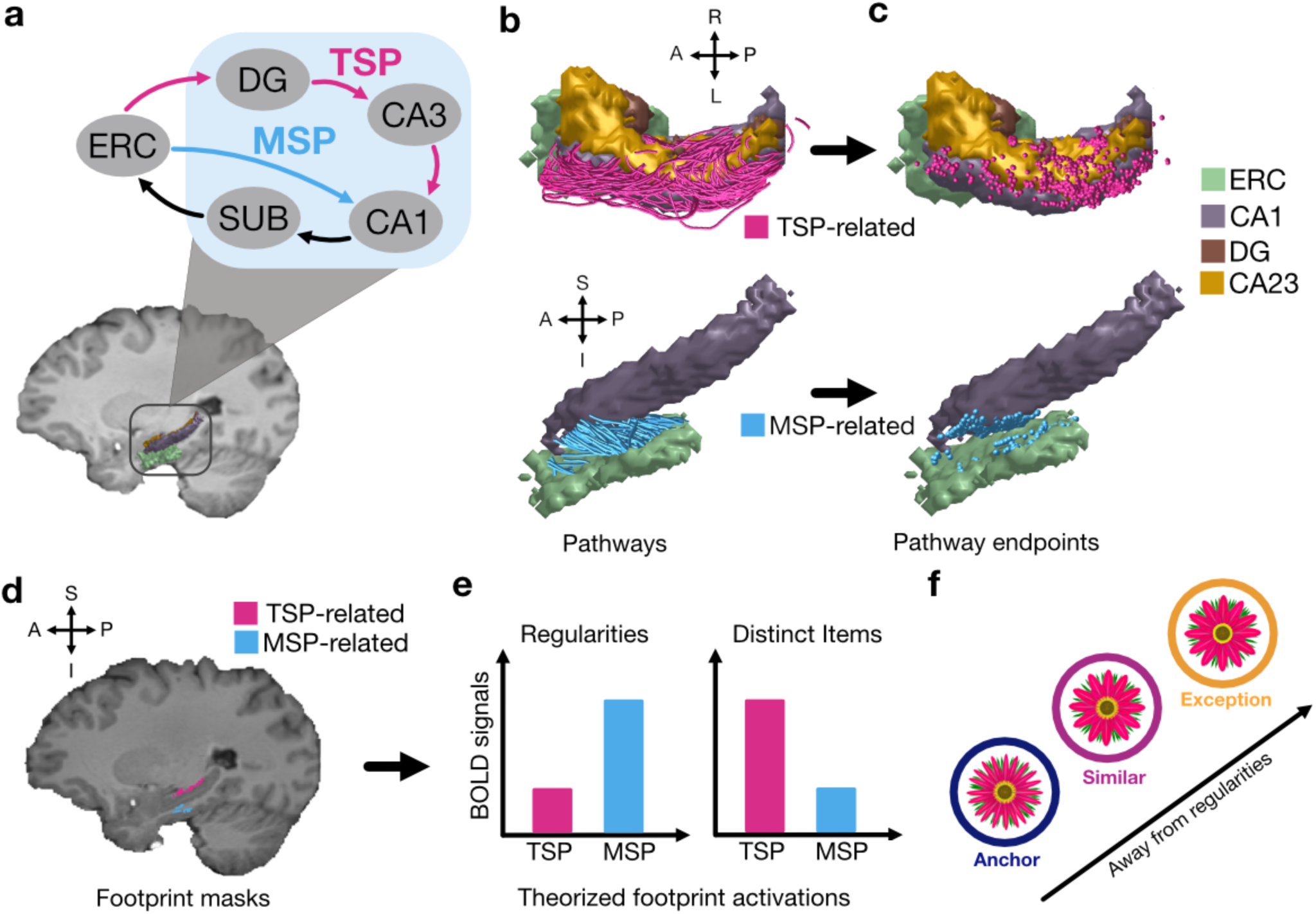
Measuring activation of hippocampal pathway footprints during concept learning. **a,** Theorized hippocampal circuitry. Trisynaptic pathway loops through ERC, DG, CA3, CA1 while monosynaptic pathway directly connects ERC to CA1. **b,** TSP and MSP related fibers are characterized using diffusion weighted imaging. **c,** End points of quantified MSP and TSP fibers. **d,** End points of TSP and MSP are mapped out using track density imaging, resulting in TSP and MSP footprint masks. **e,** BOLD signals collected during learning are masked with MSP and TSP footprint masks to measure pathway involvement. MSP is theorized to be involved in extracting regularities while TSP encodes distinct information. **f,** BOLD signals are collected during a concept learning task that includes regularities (i.e., anchors) and distinct items (i.e., similar items and exceptions). We leverage the pathway footprint approach to investigate hippocampal function in encoding regularities and distinct items during new concept learning. Concept learning is the process of grouping items as similar kinds based on their shared features (e.g., most birds have wings and fly); encoding such regularities is key to generalizing concept knowledge to novel experiences. However, natural concepts often include items that violate regularities (e.g., bats have wings and fly but are mammals, not birds). Prominent learning theories (Love et al., 2004; Nosofsky et al., 1994) posit that these concept structures are learned with adaptive encoding wherein exceptions require more specialized representations in memory structures than items that follow regularities. Notably, recent work suggests that exceptions are incorporated into knowledge after regularities have been learned (Heffernan et al., 2021; Perović et al., 2023; Y. Xie & Mack, 2024), suggesting distinct encoding dynamics early versus late in learning. The hippocampus, with its complementary operations for distinct encoding via TSP and extracting regularities via MSP (Fig. 1e), makes it an ideal candidate for exception learning, a proposal with mounting empirical support (Davis et al., 2012a, 2012b; Schlichting et al., 2021). Indeed, recent evidence suggests the strength of estimated white matter connections between TSP-related subfields is specifically associated with learning ability for exceptions (Schlichting et al., 2021). While computational models posit a functional dichotomy between hippocampal pathways for encoding regularities versus unique items (Schapiro et al., 2017; Sučević & Schapiro, 2023), their contributions are yet to be empirically validated. We bridge this crucial gap by translating a computational theory into a testable learning paradigm in humans. By leveraging a novel technique – pathway footprints – we test the specific hypothesis that MSP is preferentially engaged during the initial learning of regularities, while TSP is subsequently recruited for the acquisition of exceptions (Fig. 1f).

A notable challenge to investigate the hippocampus with functional Magnetic Resonance Imaging (fMRI) is translating theorized functions important for memory and learning such as pattern separation, differentiation, and integration into testable signatures of blood-oxygen-level-dependent (BOLD) response. For example, it has been argued that pattern separation processes in DG and CA3 can be identified by heightened BOLD response (Bakker et al., 2008). However, pattern separation creates sparse representations that, by definition, result in independent patterns of relatively fewer activated neurons (Knierim & Neunuebel, 2016; Leutgeb et al., 2007; Niibori et al., 2012), which is one way that a relative decrease in BOLD response has been interpreted (Aly & Turk-Browne, 2016). Indeed, the complex nature of cognitive function associated with increased or decreased BOLD response is one motivation for the current work. Multivariate methods that characterize the information contained within patterns of neural activation offer one solution to this confusion; compelling demonstrations of pattern separation and differentiation for overlapping experiences have been indexed by marked changes in the similarity of hippocampal activation patterns (Kim et al., 2017; Schlichting et al., 2015; Stokes et al., 2015; Wanjia et al., 2021). However, such analyses depend on robust estimates of activation patterns which present an additional empirical challenge, especially when characterizing the transient nature of learning operations that quickly unfold across a few experiences. Moreover, hippocampal pathways span multiple subfields, thus understanding their involvement in complex learning and memory necessitates an approach beyond the traditional techniques that often treat activation signatures of hippocampal subfields independently.

We propose a novel signature of hippocampal pathway engagement that characterizes univariate activation within anatomically derived regions of hippocampal subfield interconnections. We define these anatomical regions of subfield interconnection based on estimates of white matter connectivity measured with diffusion weighted imaging (Fig. 1b). Although diffusion methods offer an indirect measure of anatomy, recent diffusion studies have estimated *in vivo* connection of the human hippocampus (Dalton et al., 2022; Gumus et al., 2025; Huang et al., 2021; Karat et al., 2024; Maller et al., 2019) and demonstrated important relationships between intrinsic hippocampal connectivity and complex learning and memory behaviour (Schlichting et al., 2021; Yassa et al., 2011). Here, we uniquely leverage this approach to define “pathway footprints” based on the endpoints of estimated white matter fibers that connect subfields (Fig. 1c). These footprints serve as target locations (i.e., functional footprints) for characterizing functional interactions of subfields within MSP and TSP (Fig. 1d). Specifically, we hypothesize that the more a pathway is engaged in a learning condition, the more these regions of subfield interconnection will be activated. By constraining the functional search to anatomical signatures of subfield connections, we aim to identify the relative contribution of hippocampal pathways via univariate signal more precisely.

## 2. Results

### 2.1. Individual Variability in Exception Learning

Participants performed a feedback-based concept learning task that had a rule-plus-exception category structure (Fig. 2a) (Shepard et al., 1961). Each concept (i.e., sun vs. shade preferring flowers) included anchors, as well as similar items and exceptions that differed from their concept anchor by 1 and 2 features, respectively (Fig. 2b). These aspects of the concepts were acquired across 6 learning blocks (Fig. 2c). To test our prediction for temporal dynamics of hippocampal pathways in encoding regularities and exceptions, we separated the learning process into early (first 4 blocks) versus late (last 2 blocks) phases. This labelling was determined by identifying which block first showed performance of 50% or above for learning anchors for all participants to ensure that concept regularities were acquired. This was true in block 5 (Fig. 2c inset points; Fig. S1). At the end of learning, participants learned (Fig. 2d) anchors well (accuracy of last two learning blocks, M = 89.49%, SD = ±13.10), similar item performance was lower (M = 68.17%, SD = ±12.84), and exception learning was the lowest but also highly variable (M = 47.52%, SD = ±16.65). As such, we aimed to characterize the association between variability in learning behaviour and individual differences in neural activation related to hippocampal pathways during early and late learning.

**Fig. 2:**
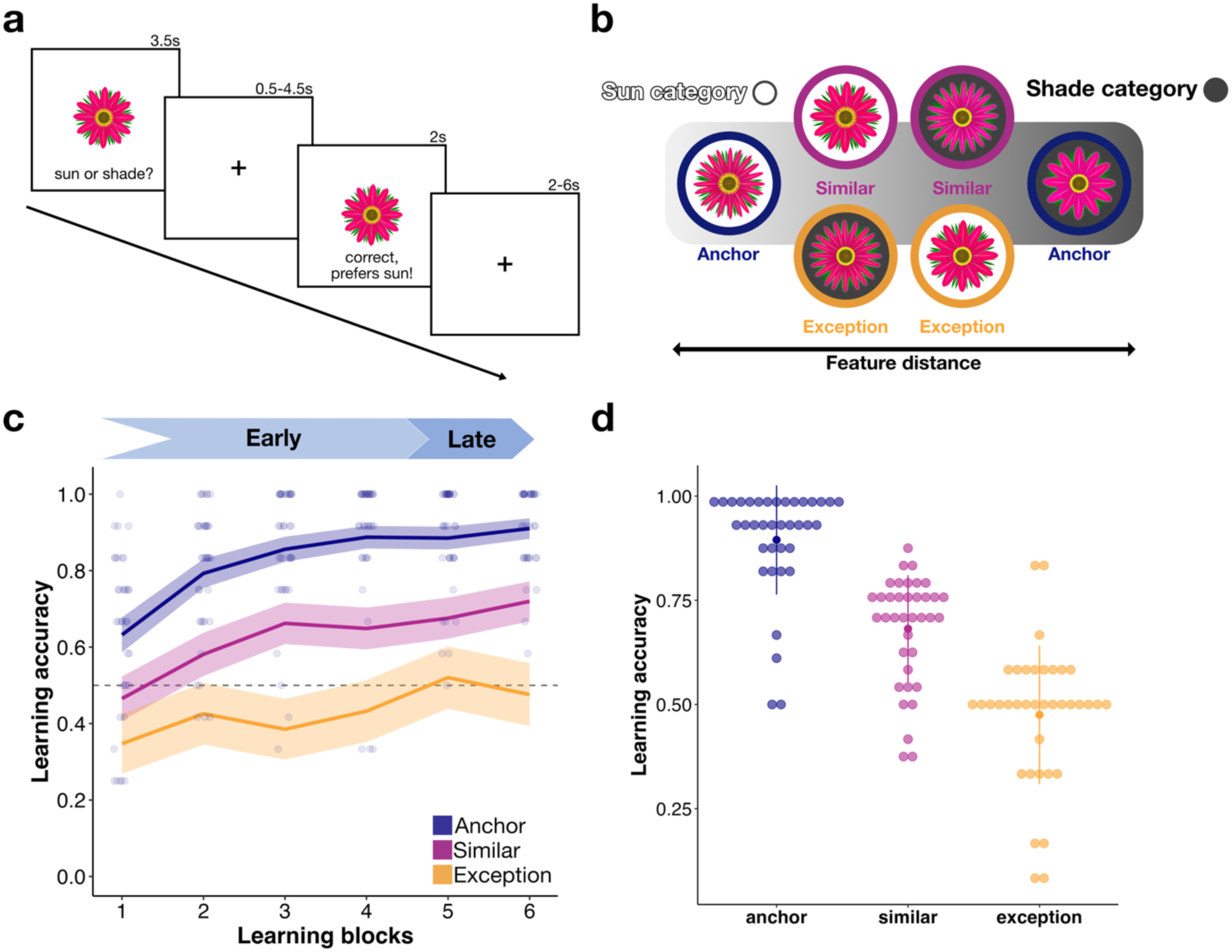
Concept learning task design and performance. **a,** Participants decided whether the presented flower on each trial preferred sun or shade and received feedback immediately. **b,** Subway plot depicting concept structures. Dark filled circles represent the shade concept while white filled circles belong to the sun concept. Similar items (pink) are 1 and exceptions (orange) are 2 features away from their concept anchors (purple). Exceptions are closer to the anchor of the other concept in feature space, making them more distinct. **c,** Learning performance across learning blocks. Points represent individual learning performances for anchors in each experimental block. The dashed line represents the chance level. All participants reached 50% or above performance in learning anchors by block 5. **d,** End of learning performance based on the last 2 blocks out of total of 6; above chance performance for anchors but individual variability for similar items and exceptions.

### 2.2. Early MSP-related Footprint Activation Relates to Regularity Learning

To characterize the relative roles of hippocampal pathways in encoding concept regularities and their exceptions early versus late learning, average activation within anatomically derived pathway footprints was calculated separately for the different stimulus types in each learning block. Mixed-effects regression was then leveraged to estimate the relationship between participant-specific indices of pathway engagement and learning accuracy during early (i.e., blocks 1-4) and late (i.e., blocks 5-6) learning. As our experimental paradigm included within-participant factors (stimulus types and learning phases) as well as continuous regressors (learning performance), a mixed-effects model was leveraged to estimate the fixed effects related to the experimental conditions while allowing for different baselines in pathway activation across participants (i.e., a random effect of intercept). Thus, this enabled us to evaluate the degree that MSP- or TSP-related footprint activation was associated with individuals’ learning of concept regularities and exceptions and if this varied throughout learning. Consistent with the hypothesized role of hippocampal pathways (Schapiro et al., 2017; Sučević & Schapiro, 2023), we found a robust significant interaction for learning performance and pathway-related end point activations, both for regularities (i.e., anchors) early in learning (β=1.60 95% CI [0.26, 2.94], p = 0.02) and exceptions late in learning (β=2.20 95% CI [0.91, 3.49], p = 0.001), suggesting a dynamic MSP and TSP engagement for encoding regularities and exceptions in learning.

Specifically, during early learning, anchor performance (Fig. 3a, first panel) was significantly associated with higher MSP-related footprint activation (β=1.74, 95% CI [0.89, 2.59], p < 0.001) but lower TSP-related footprint activation (β=-0.92, 95% CI [-1.64, -0.21], p = 0.012). Both of these effects were robust to within-participant permutation tests (p<0.01; Fig. 3a, first panel inset). There was no relationship between anchor learning and MSP-related (β=0.74, 95% CI [- 0.41, 1.89], p = 0.20) or TSP-related (β=-0.32, 95% CI [-1.11, 0.48], p = 0.43) footprint activation late in learning (Fig. 3a, second panel). Early functional MSP- and TSP-related footprints related to anchor learning are depicted on a sample participant brain (Fig. 3b); MSP-related footprint activation appears more anterior than that of TSP.

**Fig. 3:**
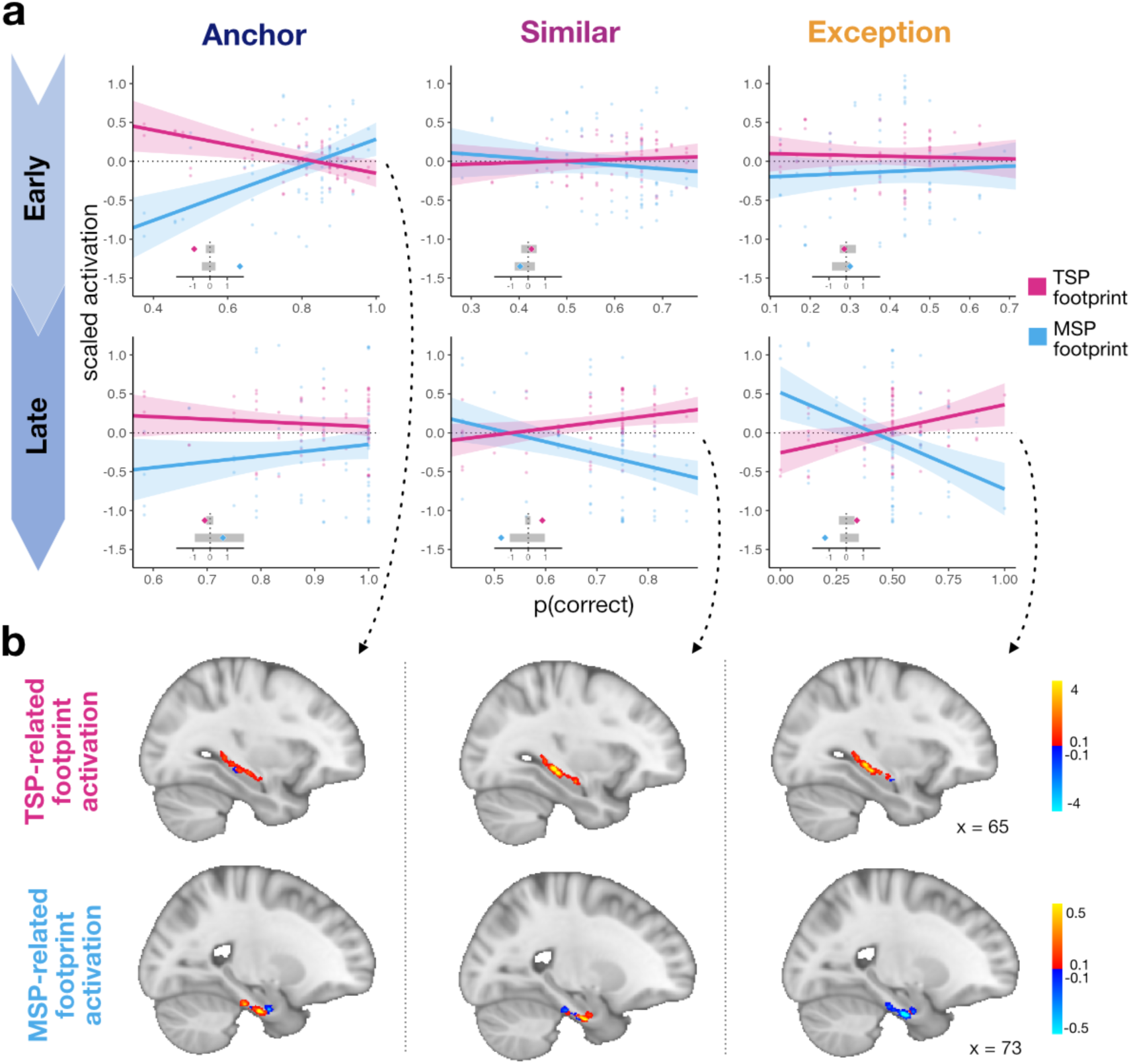
Distinct MSP- and TSP-related footprint activation in learning regularities versus similar items and exceptions. **a,** Marginal estimates of the learning accuracy effect on TSP- and MSP-related footprint activation (lines and shaded confidence bands; points represented participant-specific data) are depicted separately for early versus late learning. Insets depict permutation test results with 95% confidence interval of the null distribution (shaded grey region) and estimated coefficient of accuracy-activation effect (diamonds). Early in learning, higher MSP-but lower TSP-related footprint activation was significantly associated with better learning performance for anchors. Late in learning, higher TSP-but lower MSP-related footprint activation was correlated with learning performance for similar items and exceptions. **b,** MSP- and TSP-related footprint activation maps during early and late learning from a representative participant.

### 2.3. Late TSP-related Footprint Activation Relates to Similar Item and Exception Learning

We also evaluated our hypothesis that the engagement of TSP-related footprints late in learning relates to encoding of similar items and exceptions that are one and two features away from concept regularities (Fig. 1f). As predicted, late in learning, accuracy for similar items was significantly correlated with higher TSP-related footprint activation (β=0.82, 95% CI [0.11, 1.54], p = 0.025) and lower MSP-related footprint activation (β=-1.58, 95% CI [-2.48, -0.68], p < 0.001). These effects were robust to within-participant permutation tests (p<0.001; Fig. 3a, second panel inset). TSP-related engagement for similar items was specific to late in learning, as we found no association between learning performance for similar items and MSP-(β=-0.47 95% CI [-1.40 0.46], p = 0.32) nor TSP-related (β=0.20, 95% CI [-0.60, 0.99], p = 0.63) footprint activation early in learning (Fig 3a, second panel).

Similarly, late in learning, exception accuracy was significantly associated with higher TSP-related footprint activation (β=0.62, 95% CI [0.09, 1.15], p = 0.02) and lower MSP-related footprint activation (β=-1.24, 95% CI [-1.87, -0.61], p < 0.001; Fig. 3a, third panel). These effects were robust to random permutation tests (p<0.001; Fig. 3a, third panel inset). As hypothesized, TSP-related footprint involvement in exception learning was only observed late in learning; exception learning performance was not related to MSP-(β=0.23, 95% CI [-0.62, 1.08], p = 0.60) or TSP-related (β=-0.11, 95% CI [-0.84, 0.61], p = 0.76) footprint activations early in learning (Fig. 3a, third panel).

### 2.4. Activation of Hippocampal Subfields Does Not Fully Explain Learning Dynamics

We found robust effects linking learning performance to activation within pathway footprints; however, it may be the case that these effects are driven by subfield activation across their entire extent irrespective of the pathway constraint. To test the specificity of our findings, we investigated the relationship between the univariate activation of hippocampal subfields and ERC, and learning performance for the different stimulus types (Fig. 4a). In contrast to the pathway findings, no region showed activation that related to learning performance for any of the stimulus types (all p > 0.12) except for late activation of ERC for similar items (β=-1.30, 95% CI [-2.47, -0.12], p < 0.05; Table S1).

**Fig. 4:**
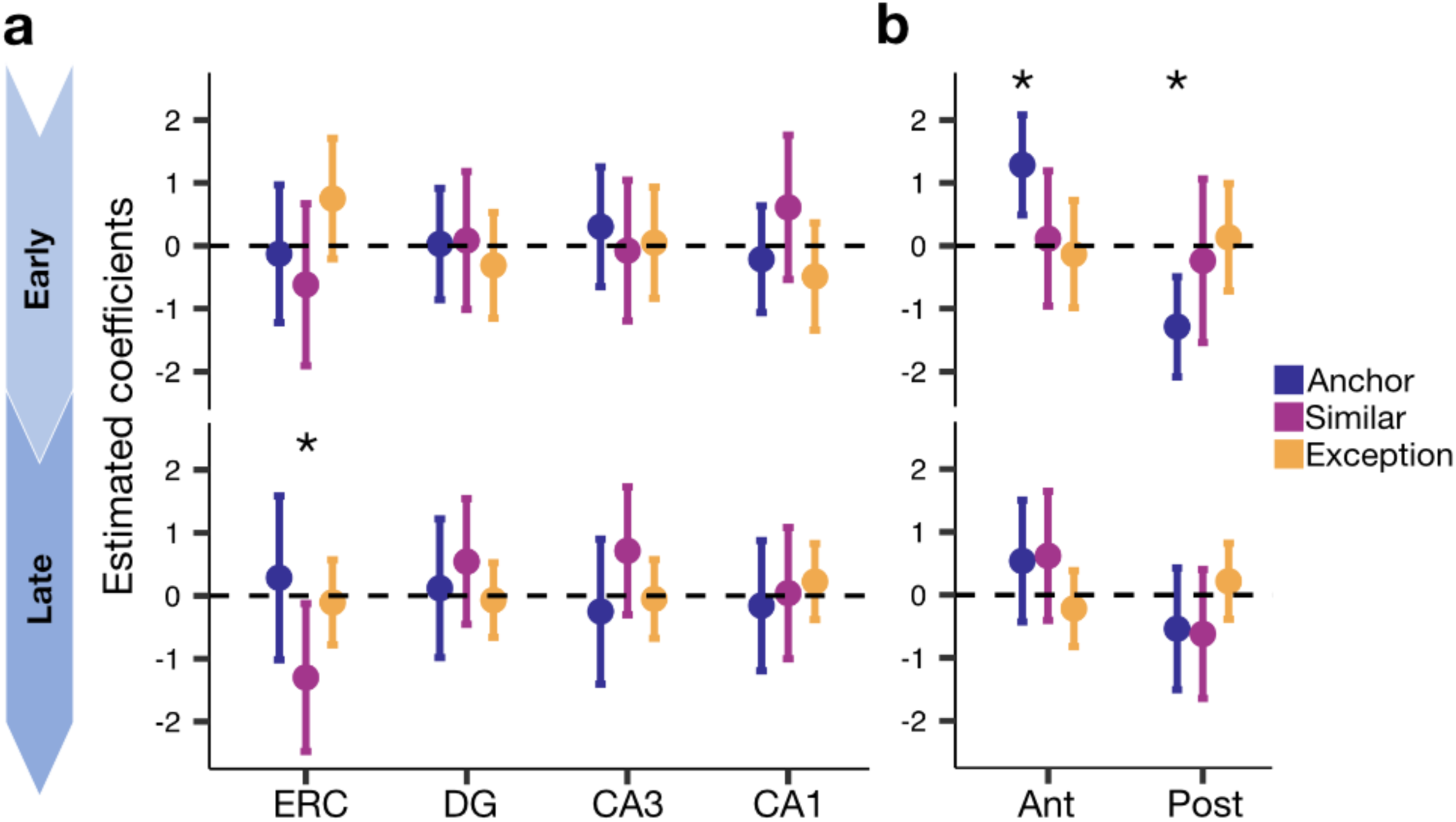
Traditional analyses of volume-based univariate activations do not fully capture learning dynamics. **a,** Deactivation of ERC early in learning was related to similar performance. **b,** High anterior (ant) but low posterior (post) hippocampus activation early in learning was correlated with anchor learning.

### 2.5. Early Activation of Anterior Hippocampus Relates to Regularity Learning

Given that ERC is located closer to the anterior extent of the hippocampus, the pathway footprints necessarily differ in their coverage along the hippocampal long axis (i.e., MSP-related footprint is relatively more anterior, TSP-related footprint is more posterior). It is well established that key functional differences exist along the hippocampal anterior-posterior axis (Poppenk et al., 2013; Strange et al., 2014) which opens the possibility that the footprint effects we observed are due to distinct functional engagement of anterior versus posterior hippocampus rather than the regions’ relationship to pathways. To test this alternative account, we investigated whether the activation within anterior and posterior hippocampus is linked to learning. We found that early in learning, higher activation of anterior (β=1.28, 95% CI [0.49, 2.08], p < 0.05) but reduced activation of posterior hippocampus (β=-1.28, 95% CI [-2.08, -0.49], p < 0.05) were significantly associated with anchor performance (Fig. 4b). However, no other effects were observed (all p>0.23; Table S2). These control analyses suggest that hippocampal pathway interconnections, not whole subfields nor anterior/posterior divisions, provide a meaningful target for linking neural engagement to learning.

## 3. Discussion

We introduce a novel technique, pathway footprints, that leverages diffusion-weighted imaging and task-based fMRI to investigate the involvement of hippocampal pathways in learning. Specifically, we estimated white matter streamlines connecting hippocampal subfields and ERC and constrained analysis of univariate activation to only regions that included endpoints of these connections (i.e., footprints). By sorting the footprints between ERC and hippocampal subfields according to the monosynaptic (ERC-CA1) or trisynaptic (ERC-DG-CA3-CA1) pathways, we tested the hypothesis that hippocampal pathways distinctly support learning regularities and distinct items. Indeed, consistent with neurobiological and computational frameworks (McClelland et al., 1995; O’Reilly & Rudy, 2001; Schapiro et al., 2017; Sučević & Schapiro, 2023), we find that MSP-related footprint activation is associated with early learning of regularities across experiences (i.e., anchors). On the other hand, late in learning, TSP-related footprint activation supports the encoding of distinct items (i.e., similar and exception items) that must be reconciled with concept regularities. This is in line with prior work that demonstrate that exception learning is facilitated after concept regularities have been learned (Heffernan et al., 2021; Perović et al., 2023; Y. Xie & Mack, 2024). Notably, these findings were specific to the pathway footprint approach; conventional approaches to parcellating the hippocampus into subfields and anterior and posterior extents revealed no such relationship to learning. As such, we propose that the pathway footprint method offers a more precise estimate of neural signatures related to the relative engagement of hippocampal pathways. By focusing on the regions of interconnection between MSP and TSP, we provide unique empirical support for the dynamic role of the hippocampus during complex learning.

Our findings suggest that the theorized complimentary functions of the hippocampus are implicated at different time points during learning, perhaps due to the quickly evolving demands of the learning task we employed. We found that MSP-related footprint activation early in learning was associated with learning concept anchors, the stimuli that represent concept regularities, consistent with the hippocampal role in generalization. Recent work has established the role of the hippocampus in building categories and generalization of learned categories (Theves et al., 2021; Zeithamova et al., 2012; Zeithamova & Bowman, 2020). Computationally, generalization is thought to arise from pattern completion and integration mechanisms (Schapiro et al., 2012, 2017; Sučević & Schapiro, 2023). These memory functions that act to incorporate new information into existing memory are often supported by CA1 (Duncan et al., 2012; Larkin et al., 2014; Schlichting et al., 2014; Wang & Morris, 2010), a key region in MSP. However, we found no link between the univariate activation of CA1 and learning. Rather, only by grounding the search for pathway function in its anatomical structure, were we able to implicate MSP in building concept regularities. That this effect occurred early in learning suggests MSP may uniquely build concept regularities that are leveraged by TSP for later encoding of distinct items.

We also found that once the concept regularities are established, TSP is engaged in learning of similar and exception items forming flexible concepts, in line with its theorized role (Schapiro et al., 2017; Sučević & Schapiro, 2023). Exceptions in our task are unique as they are visually more similar to the regularities of the other concept than their own, presumably demanding the formation of separate neural patterns. Although similar items are not as distinct as exceptions, their unique position between exceptions and anchors in feature space necessitates their reconciliation with concept regularities. Both neural and computational modelling work suggest that encoding exceptions require distinct representations in memory, especially within the hippocampus (Davis et al., 2012a, 2012b; Love et al., 2004). Differentiation of distinct elements is supported by DG and CA3 activation (Bakker et al., 2008; Wanjia et al., 2021) as well as the white matter connections between DG and CA3 (Yassa et al., 2011), and CA3 and CA1 (Schlichting et al., 2021). Yet, CA3 presents opposing functions with some evidence for its involvement in pattern integration as well (Deuker et al., 2014). The role of TSP in learning emerges from the multiple hippocampal subfields that it spans, demanding an approach beyond the traditional techniques that consider univariate activation of subfields separately. Instead, by leveraging both diffusion-weighted imaging and task-based fMRI, as we have done here, the pathway footprint method provides a comprehensive investigation into hippocampal pathways specific to the activation of their interconnections within the subfields. Our findings suggest that in reconciling similar and exception items with already formed concept regularities later in learning, TSP shows elevated engagement, the degree to which is associated with learning success.

One potential confound with the pathway footprint approach is that the footprints may simply reflect the univariate signals from hippocampal subfields. However, examining this more traditional approach to characterizing subfield involvement revealed limited links to learning. Specifically, only ERC deactivation late in learning was related to similar item performance. ERC often exhibits neural adaptation to previously active representations such as grid coding learned in one environment could generalize to a novel environment (Doeller et al., 2010; Mark et al., 2024). As similar items highly resemble the concept anchors (i.e., only one feature away from them), ERC involvement may reflect generalization efforts from concept regularities to similar items late in learning. However, this interpretation is muddied by the absence of any other relationship between hippocampal subfield activations and learning. Similarly, another concern may arise from the differential locations of MSP and TSP along the long axis of the hippocampus. Subfields are located with an anatomical gradient such that CA2/3 and CA1 are located more anteriorly, and DG is more posterior, contributing to functional differences along anterior-posterior extent (Hrybouski et al., 2019; Malykhin et al., 2010; Poppenk et al., 2013). Nevertheless, separating the hippocampus into anterior and posterior extents did not explain the pathway footprint effects. Interestingly, we did find that learning anchors was related to early anterior hippocampus activation and posterior hippocampus deactivation. This is consistent with long-axis specialization of the hippocampus in similar learning paradigms: anterior hippocampus holds generalized concepts and posterior hippocampus forms more item-specific representations (Frank et al., 2019; Mack et al., 2016; Zeithamova & Bowman, 2020). It might be necessary to encode regularities early in learning in line with our pathway footprint results but, meanwhile, suppress the regions that relate to learning of distinct items. Often, deactivation might indicate the demand for sparse representations of distinct items (Aly & Turk-Browne, 2016; Schapiro et al., 2017; Sherman et al., 2023; Sherman & Turk-Browne, 2020). However, there was no neural link to learning similar items or exceptions, resulting in an incomplete picture of hippocampus long-axis involvement in learning. Overall, these control analyses highlight the precision of pathway footprint method in measuring neural signatures of learning, introducing a promising avenue for investigations of hippocampal pathway contributions to learning as well as numerous cognitive functions.

The pathway footprint approach offers an insightful window into the role of hippocampal pathway in learning by incorporating structural and functional neuroimaging modalities. Information arrives to the hippocampus from ERC and is transferred from one subfield to another one via the hippocampal pathways. Theoretically, the end points of the pathways (i.e., footprints) represent computationally specific regions of hippocampal subfields: waypoints where carried information is received and unpacked. Subsequent processing of this information may engage the rest of the subfield in functionally distinct ways before being passed onto the next stop in the pathway. Our findings suggest that this information flow through the pathways engages different functions depending on the context of a learning experience. For example, when extracting regularities from similar experiences (i.e., learning concept anchors), MSP-related footprint activation was related to successful regularity learning. In conditions requiring pattern separation and differentiation (i.e., learning exceptions to concept regularities), we observed heightened activation from TSP-related footprints that related to learning outcome. Our results provide compelling evidence that the footprint method indexes the relative engagement of MSP- and TSP-related processes in learning.

In conclusion, we revealed the dynamic involvement of hippocampal pathways in learning; MSP extracts concept regularities early in learning and TSP encodes distinct elements late in learning. This novel empirical evidence in humans for prominent theories and computational models (Schapiro et al., 2017; Sučević & Schapiro, 2023) was only possible through an analytic innovation, the pathway footprint method. Investigating hippocampal engagement with anatomical constraints of pathways offers a promising multimodal method for characterizing the role of the hippocampus in learning, memory and cognition broadly.

## 4. Methods

### 4.1. Participants

Portions of this work are based on a previously reported dataset (Schlichting et al., 2021). The study was approved by the University of Toronto Research Ethics Board. Participants were recruited from the University of Toronto community (N=43, 24 female, 19-33 years old, mean 23.5 years old). Inclusion criteria included being right-handed, having normal or corrected-to-normal vision, and having no diagnosis of neurological or psychological disorders. Written consent was collected from each participant before the study, and they were compensated $20/hour for their participation. 6 participants were excluded from the dataset because of failing to follow the experiment instructions (N=2), technical issues (N=3), and failure to estimate white matter fibers (N=1).

### 4.2. Concept Learning Task

Participants performed a concept learning task in the MRI scanner. Visual stimuli were back projected on a screen which was visible to the participants on a mirror attached to the MR head coil. The task consisted of learning to categorize cartoon flowers composed of three binary-valued feature dimensions: outer petal shape, inner petal shape and colour of the flower center. Participants were instructed to learn how to correctly categorize the flowers as preferring sun versus shade based on a combination of the flower dimension values (Fig. 2b). The categories were based on a rule-plus-exception structure (Shepard et al., 1961) with two concept “anchors”, two “similar” items per concept that mismatched their concept anchor by 1 feature, and one “exception” per concept that mismatched their respective anchor by 2 features (Fig. 2b). In fact, exceptions were visually more similar to the anchor of the opposing concept than their own. On each trial, participants were visually presented with a flower for 3.5s during which they made a concept decision by pressing one of two buttons on a MR-compatible button box (Fig. 2a). After a fixation screen of variable duration, corrective feedback including the flower image and its correct concept was displayed for 2s. To further support the rule-plus-exception category structure by making similar and exception items rare events, concept anchors were shown more frequently (each 6 trials/run) than similar items and exceptions (each 2 trials/run). One learning run consisted of 24 trials and the trial order was randomized. The entire experiment included 6 runs (144 trials total) and lasted approximately 40 minutes.

### 4.3. MRI Acquisition

Whole-brain imaging data were acquired on a 3T Siemens Prisma system at the Toronto Neuroimaging Facility housed at the University of Toronto. A high-resolution T1-weighted MPRAGE structural volume (TR = 2s, TE = 2.4ms, flip angle = 9°, FOV = 256mm, matrix = 256×256, 1mm iso-voxels) was acquired for co-registration and parcellation. Two oblique coronal T2-weighted structural images were acquired perpendicular to the main axis of the hippocampus (TR = 4s, TE = 66ms, matrix = 512×512, 0.43×0.43mm in-plane resolution, 2mm thru-plane resolution, 40 slices, no gap). Functional images were also acquired using a T2*-weighted EPI pulse sequence during the concept learning task (TR = 2 s, TE = 30 ms, flip angle = 90°, FOV = 220 mm, matrix = 130 × 130, slice thickness = 1.7 mm, multiband factor = 4) allowing for whole brain coverage with 1.7-mm isotropic voxels. Diffusion weighted images were also collected in two separate scans one before and one after the learning task (axial echo-planar imaging, GRAPPA=2, multiband factor=2, b-value=1000, TR=4s, TE=70ms, 64 directions, FOV=220mm, 2mm-iso voxels, 70 slices, acquisition time=298s; sequence also included 4 repetitions of b-value=0 images). An additional two reverse-phase encoded b-value=0 images were collected for distortion correction.

### 4.4. Regions of Interests

Hippocampal subfields and ERC were segmented with the Automatic Segmentation of Hippocampal Subfields (ASHS) (Yushkevich et al., 2014) tool and the Princeton Young Adult 3T ASHS atlas (Hindy et al., 2016) using T2 weighted images. This approach generated participant-specific regions of interests including ERC and hippocampal subfields DG, CA1, combined CA2 and CA3 (labelled as CA3 here), and subiculum in both left and right hemispheres. These regions of interests (ROIs) were registered to participants’ T1 anatomical space using the linear transformations provided by ASHS. We also segmented anterior and posterior hippocampus with ASHS using the ASHS Penn Memory Center T1-Only Atlas (PMC-T1) (L. Xie et al., 2016), using T1-weighted images of each participant. All ROIs were registered to the BOLD space of participant specific runs, using Advanced Normalization Tools (ANTs) (Avants et al., 2011).

### 4.5. Construction of Hippocampal Connectomes

We first created participant specific whole brain tractographies using MRtrix3 (Tournier et al., 2019), which allows characterization of fiber populations within each voxel (i.e., fixel) through constrained spherical deconvolution and produces a precise estimation of crossing fiber orientation (Tournier et al., 2012). Based on individual whole brain tractographies, we used the segmentations of hippocampal subfields and ERC to create hippocampal connectivity matrices, where the value in each entry represents the streamline density between pairs of regions. Using these connectomes, we then reconstructed the fibers specific to hippocampal circuitry. We further isolated the fibers between each pair of ROIs, using *connectome2tck* (Tournier et al., 2019). The quantified fibers between ERC and CA1 were labelled as MSP (Fig. 1a-b). On the other hand, TSP included the fibers between DG-CA2/3 and CA2/3-CA1 (Fig. 1a-b). The connections between ERC-DG were not included within the analyses due to the difficulty in isolating these specific fibers.

### 4.6. Hippocampal Pathway Specific Footprints

Isolated fibers consist of 2 sets of endpoints. For instance, the MSP specific fibers have ends both on ERC and CA1 (Fig. 1c). The tract density of these end points for each isolated pathway were saved as separate images (i.e. footprint masks; Fig. 1d), using *tckmap* (Tournier et al., 2019). These density maps are used as the regions of interest (ROIs), where assessed activation would represent the pathways endpoint engagement. This resulted in separate footprint masks for each MSP (i.e., ERC-CA1) and TSP (DG-CA2/3 and CA2/3-CA1) connections (Fig. 1d, Fig. S2a-c). Each voxel in these footprint masks represent the density of the end points, or more generally the density of the streamlines landing on this voxel (Fig. 1d, Fig. S2a-c). There was no overlap between the fiber endpoints on CA1 that receives from ERC (i.e., more inferior of CA1) and CA2/3 (i.e., more superior) (Fig. S2d). The footprints were then linearly registered to the BOLD image from each run at the participant level, using Advanced Normalization Tools (ANTs) (Avants et al., 2011). We then used the footprints to mask the BOLD signals for each stimulus type, in run and in each participant (Fig. 1d). The measured activations are specific to pathway footprints (i.e., specific voxel locations) on each ROI. Thus, by masking the BOLD images with footprints instead of the entire ROI volume, signal in each voxel is weighted by streamline density, allowing us to constrain the search for pathway function by anatomy.

### 4.7. Preprocessing fMRI Data

Functional volumes were preprocessed with fmripep version 20.2.1 (Esteban et al., 2019; Esteban, Markiewicz, DuPre, et al., 2020), which is based on Nipype 1.5.1 (Esteban, Markiewicz, Burns, et al., 2020; Gorgolewski et al., 2011). For each of the BOLD runs found per subject (across all task blocks), the following preprocessing was performed. First, a reference volume and its skull-stripped version were generated using a custom methodology of fMRIPrep. A B0-nonuniformity map (or fieldmap) was estimated based on a phase-difference map calculated with a dual-echo GRE (gradient-recall echo) sequence, processed with a custom workflow of SDCFlows inspired by the epidewarp.fsl script and further improvements in Human Connectome Project (HCP) Pipelines (Glasser et al., 2013). The fieldmap was then co-registered to the target EPI (echo-planar imaging) reference run and converted to a displacements field map (amenable to registration tools such as ANTs) with FSL’s fugue and other SDCflows tools.

Based on the estimated susceptibility distortion, a corrected EPI (echo-planar imaging) reference was calculated for a more accurate co-registration with the anatomical reference. The BOLD reference was then co-registered to the T1w reference using flirt (FSL 5.0.9) (Jenkinson & Smith, 2001) with the boundary-based registration (Greve & Fischl, 2009) cost-function. Co-registration was configured with nine degrees of freedom to account for distortions remaining in the BOLD reference. Head-motion parameters with respect to the BOLD reference (transformation matrices, and six corresponding rotation and translation parameters) are estimated before any spatiotemporal filtering using mcflirt (FSL 5.0.9) (Jenkinson et al., 2002). The BOLD time-series (including slice-timing correction when applied) were resampled onto their original, native space by applying a single, composite transform to correct for head-motion and susceptibility distortions. These resampled BOLD time-series will be referred to as preprocessed BOLD in original space, or just preprocessed BOLD. The BOLD time-series were resampled into standard space, generating a preprocessed BOLD run in MNI152NLin2009cAsym space. First, a reference volume and its skull-stripped version were generated using a custom methodology of fMRIPrep. Several confounding time-series were calculated based on the preprocessed BOLD: framewise displacement (FD), DVARS and three region-wise global signals. FD was computed using two formulations following Power (absolute sum of relative motions) (Power et al., 2014) and Jenkinson (relative root mean square displacement between affines) (Jenkinson et al., 2002) FD and DVARS are calculated for each functional run, both using their implementations in Nipype (following the definitions by Power et al.,) (Power et al., 2014). The three global signals are extracted within the CSF, the WM, and the whole-brain masks. Additionally, a set of physiological regressors were extracted to allow for component-based noise correction (CompCor) (Behzadi et al., 2007). Principal components are estimated after high-pass filtering the preprocessed BOLD time-series (using a discrete cosine filter with 128s cut-off) for the two CompCor variants: temporal (tCompCor) and anatomical (aCompCor). tCompCor components are then calculated from the top 2% variable voxels within the brain mask. For aCompCor, three probabilistic masks (CSF, WM and combined CSF+WM) are generated in anatomical space. The implementation differs from that of Behzadi et al. in that instead of eroding the masks by 2 pixels on BOLD space, the aCompCor masks are subtracted a mask of pixels that likely contain a volume fraction of GM. This mask is obtained by thresholding the corresponding partial volume map at 0.05, and it ensures components are not extracted from voxels containing a minimal fraction of GM. Finally, these masks are resampled into BOLD space and binarized by thresholding at 0.99 (as in the original implementation).

Components are also calculated separately within the WM and CSF masks. For each CompCor decomposition, the k components with the largest singular values are retained, such that the retained components’ time series are sufficient to explain 50 percent of variance across the nuisance mask (CSF, WM, combined, or temporal). The remaining components are dropped from consideration. The head-motion estimates calculated in the correction step were also placed within the corresponding confounds file. The confound time series derived from head motion estimates and global signals were expanded with the inclusion of temporal derivatives and quadratic terms for each (Satterthwaite et al., 2013). Frames that exceeded a threshold of 0.5 mm FD or 1.5 standardised DVARS were annotated as motion outliers. All resamplings can be performed with a single interpolation step by composing all the pertinent transformations (i.e. head-motion transform matrices, susceptibility distortion correction when available, and co-registrations to anatomical and output spaces). Gridded (volumetric) resamplings were performed using antsApplyTransforms (ANTs), configured with Lanczos interpolation to minimize the smoothing effects of other kernels (Lanczos, 1964). Non-gridded (surface) resamplings were performed using mri_vol2surf (FreeSurfer).

### 4.8. Estimating Beta Series for Each Stimulus Type

Beta series of each stimulus type (i.e., anchor, similar and exception) were estimated from the preprocessed whole brain fMRI images, using the Ordinary Least Squares (OLS) approach (Holmes & Friston, 1998). Beta series were estimated for each stimulus type both for stimulus and feedback presentation. Events only related to stimulus presentations were used in the analyses. Confound regressors in the beta estimations included global signals, framewise displacement, six motion parameters (three degrees of translation and rotation) (as well as first six components of anatomical CompCor) (Behzadi et al., 2007). This regression analysis results in statistical t-maps for each stimulus type in each run in each participant. To measure the activation of the MSP- and TSP-related footprints, we masked the estimated t-maps for each stimulus type in each run with individual specific MSP- and TSP-related footprint masks. For our control analyses, these t-maps were masked with hippocampal subregions, ERC and anterior/posterior hippocampus, resulting in traditional volume based univariate activations.

### 4.9. Relating Activation of Hippocampal Footprints to Learning

All analyses were completed with R statistical software, version 4.2.1 (R Core Team, 2021). We fitted 3 separate linear mixed effects model (*lme4* version 1.1-30, *lmerTest* version 3.1-3) (Bates et al., 2015; Kuznetsova et al., 2017), that is estimated using REML and optimx optimizer, to predict the early and late MSP and TSP footprint activations with early and late learning accuracies for each stimulus type; anchor, similar item and exception. The fixed effects were an interaction of learning accuracy for the respective stimulus type, early/late phase of learning and pathway name (i.e., MSP and TSP) to predict their footprint activations. Each model also accounted for individual differences with 2 random effects: one with an intercept for each early/late phase and MSP-/TSP-related footprints at the individual level, and another intercept for each of the 2 diffusion images for a participant.

### 4.10. Bootstrap Analyses of the Link between Pathway Footprints and Learning

To assess the reliability of these complex models above, we conducted bootstrap sampling analyses. In each iteration, participant data was randomly sampled with replacement and then fit to each linear mixed effects model described above. This procedure was repeated 10,000 times for each model. We report p-value statistics based on the distributions of the coefficients of the models which represent the slopes of relationship between activation of pathway footprints and learning performance. Reported p-values correspond to the proportion of resampled coefficients that were more extreme than the observed effect.

### 4.11. Relating Univariate Activations to Learning

For our control analyses, we leveraged the univariate activations of hippocampal subfields as well as long-axis segmentations (i.e., anterior and posterior hippocampus). Much of the published work that uses multivariate techniques include a univariate control analysis with the targeted ROIs (e.g., subfields, subregions along the long axis, whole hippocampus) often finding limited effects. It is worth noting that these univariate analyses often serve as a control for the multivariate findings to ensure that effects of pattern classification or similarity are based on differences in neural patterns and not due simply to univariate differences between conditions.

We aimed to compare our footprint activations to a similar type of univariate analysis commonly used in the literature. An important validation of the footprint method is to demonstrate that any univariate effects we discovered in the pathway ROIs were specific to the pathway endpoints and not simply an effect observed across the whole subfield. For example, if we found that activation in CA3 as a whole was associated with learning performance for exceptions but not anchors, it would be likely that the TSP-footprint related mask, which includes part of CA3, would show a similar effect. Thus, this would not represent a pathway-specific finding. To ensure that an effect was localized to the anatomically defined pathway ROIs, we analyzed activation within each subfield in the same way as the pathway mask analysis to confirm that the central findings were specific to the pathways. Moreover, finding null effects in the univariate activations of the hippocampal subfields and anterior/posterior hippocampus but significant effects in pathway footprints, which are distinct regions contained within the same subfield, would provide theoretically appealing support for the novel pathway footprint method.

For our control analyses, we constructed separate linear mixed effects models to investigate the link between learning behaviour and univariate activations of; 1) hippocampal subregions and ERC, 2) anterior and posterior hippocampus. In each of these analyses, we setup 3 separate mixed effects models for each stimulus type. To predict the univariate activations, each model included a fixed effect that was modelled as the interaction of the learning accuracy for the respected stimulus, early/late phase of learning and the name of the regions of interests (i.e., hippocampal subfields/ERC or anterior/posterior hippocampus). We accounted for individual differences with random effects that fit an individual intercept for each learning phase (i.e., early versus late) and regions of interests.

## 5. Data Availability

Upon publication, the data used in this project will be publicly available at an Open Science Framework (OSF) repository.

## 6. Acknowledgements

This project was funded by Natural Sciences and Engineering Research Council (NSERC) Discovery Grants (RGPIN-2017-06753 and RGPIN-2024-0588 to MLM), Canadian Institute of Health Research (CIHR) Grant (PJT-178337 to MLM), Brain Canada Foundation Grant (to MLM), and Vanier Canada Graduate Scholarship provided by NSERC (to MG).

## 7. Author Contributions

M.G. and M.L.M. designed the study and conducted the analyses. M.G. collected the data and wrote the initial manuscript. M.L.M. supervised the project and directed project progress. Both authors edited the manuscript.

## 8. Competing Interests Statement

M.G. and M.L.M. declare no conflict of interests.

## Notes

**Conflict of interest:** The authors, MG and MLM, declare no conflict of interests.

### Competing Interest Statement

The authors have declared no competing interest.

