## Supplemental Files for "Distinct contributions of hippocampal pathways in learning regularities and exceptions revealed by functional footprints"

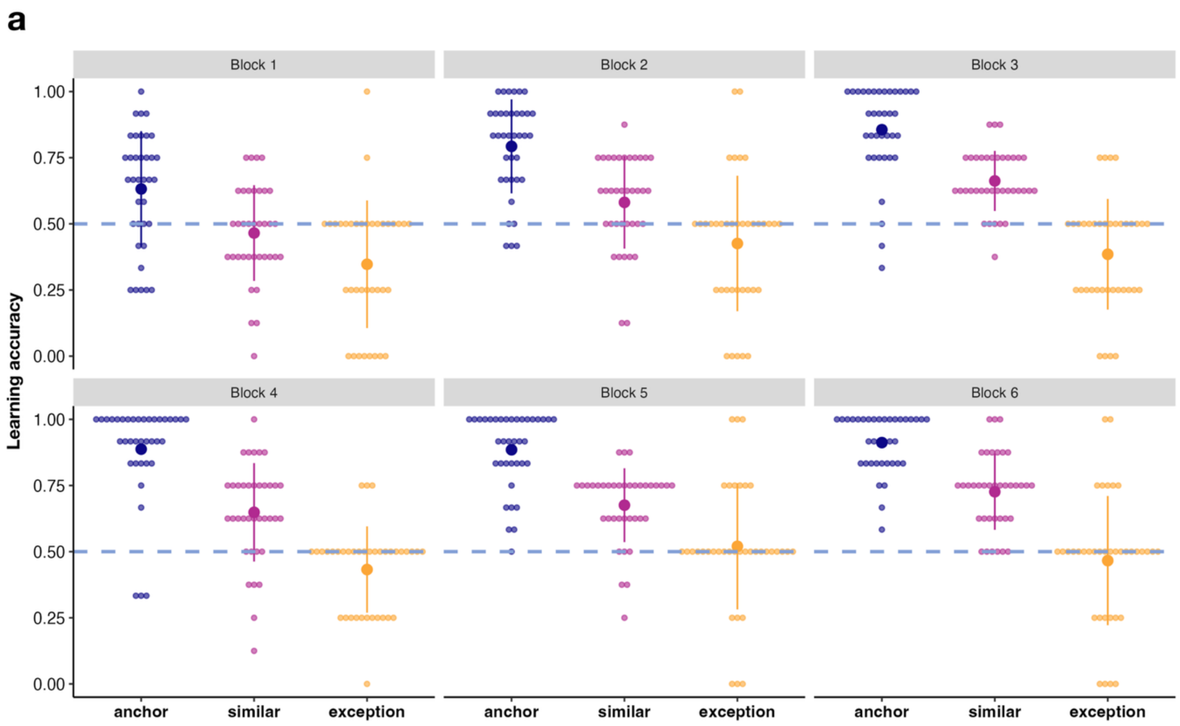


**Fig. S1. Learning performance across learning blocks.** Dots represent participants, depicting their performance for anchors, similar items and exceptions across all 6 experimental blocks.

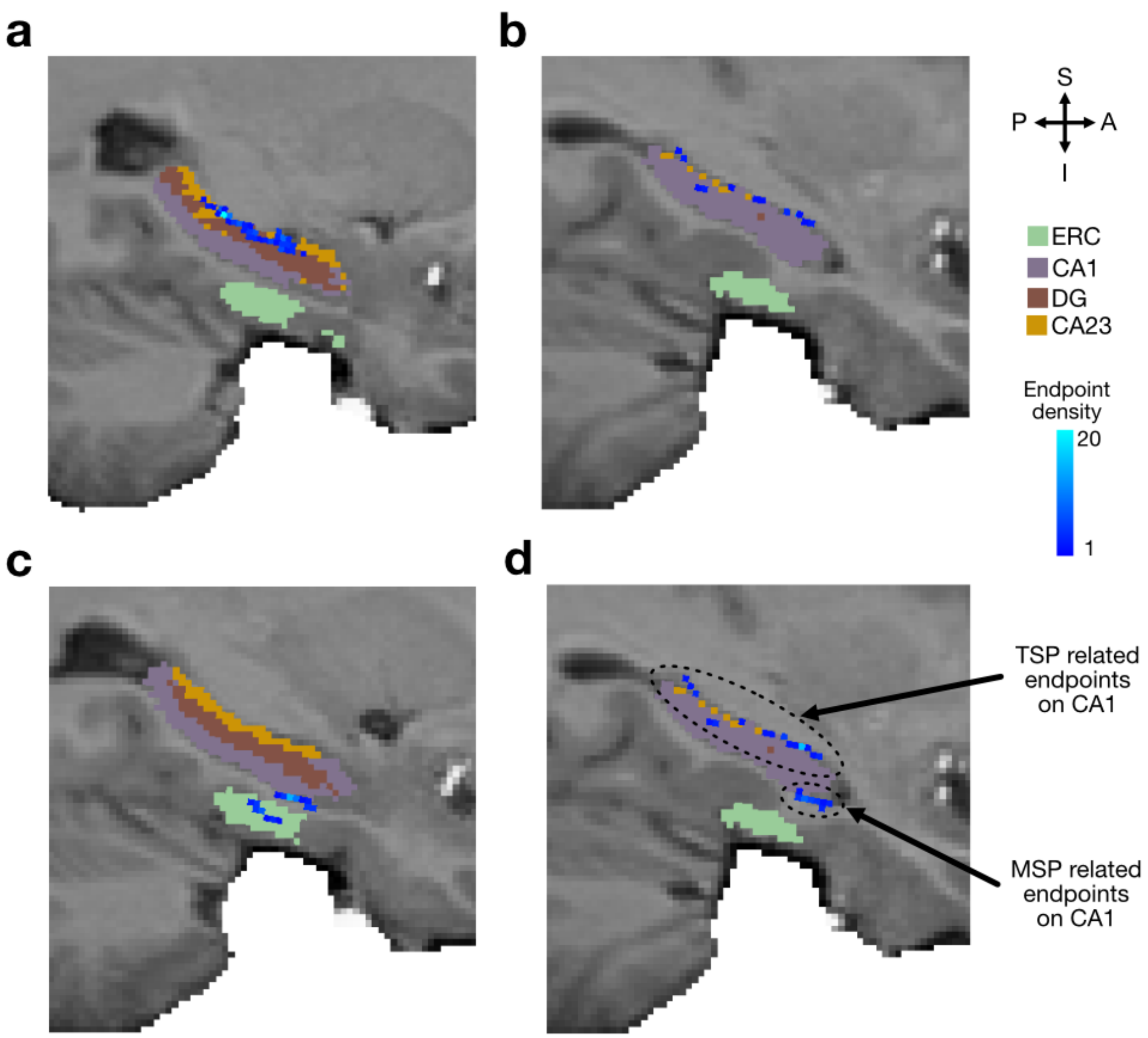


**Fig. S2. Creating hippocampal pathway specific footprints. a,** TSP-related footprints are specific to DG and CA2/3. **b,** TSP-related footprints are specific to CA2/3 and CA1. **c,** MSP-related footprints include pathway end points within ERC and CA1. **d,** Fibers emerging from ERC (i.e., MSP) versus CA2/3 (i.e., TSP) have no overlapping end points on CA1, resulting in distinct footprints for MSP and TSP-related pathways.

**Table S1. Traditional analyses of univariate activation of hippocampal subfields do not fully capture learning dynamics.**

| Learning | Stimulus type | Hippocampal subfields | Coefficients | P-values | Lower 95% | Higher 95% |
|---|---|---|---|---|---|---|
| Early | Anchor | ERC | -0.13 | 0.82 | -1.23 | 0.97 |
| | | DG | 0.03 | 0.94 | -0.85 | 0.91 |
| | | CA2/3 | 0.30 | 0.53 | -0.65 | 1.25 |
| | | CA1 | -0.21 | 0.62 | -1.06 | 0.63 |
| | Similar | ERC | -0.62 | 0.34 | -1.91 | 0.67 |
| | | DG | 0.09 | 0.88 | -1.01 | 1.18 |
| | | CA2/3 | -0.08 | 0.89 | -1.19 | 1.04 |
| | | CA1 | 0.61 | 0.29 | -0.54 | 1.76 |
| | Exception | ERC | 0.75 | 0.12 | -0.21 | 1.71 |
| | | DG | -0.31 | 0.46 | -1.15 | 0.53 |
| | | CA2/3 | 0.05 | 0.91 | -0.84 | 0.93 |
| | | CA1 | -0.49 | 0.26 | -1.34 | 0.37 |
| Late | Anchor | ERC | 0.29 | 0.66 | -1.02 | 1.59 |
| | | DG | 0.12 | 0.83 | -0.98 | 1.22 |
| | | CA2/3 | -0.25 | 0.67 | -1.40 | 0.90 |
| | | CA1 | -0.16 | 0.76 | -1.19 | 0.88 |
| | Similar | **ERC*** | **-1.30** | **0.03** | **-2.47** | **-0.12** |
| | | DG | 0.54 | 0.28 | -0.46 | 1.54 |
| | | CA2/3 | 0.71 | 0.17 | -0.30 | 1.73 |
| | | CA1 | 0.04 | 0.94 | -1.00 | 1.09 |
| | Exception | ERC | -0.11 | 0.76 | -0.78 | 0.57 |
| | | DG | -0.07 | 0.82 | -0.66 | 0.52 |
| | | CA2/3 | -0.05 | 0.87 | -0.68 | 0.57 |
| | | CA1 | 0.23 | 0.46 | -0.38 | 0.83 |

*Note.* * $p<0.05$

**Table S2. Traditional analyses of univariate activations of anterior and posterior hippocampus do not fully capture the learning dynamics.**

| Learning | Stimulus type | Hippocampal long axis | Coefficients | P-values | Lower 95% | Higher 95% |
|---|---|---|---|---|---|---|
| Early | Anchor | **Anterior*** | **1.28** | **0.002** | **0.49** | **2.08** |
| | | **Posterior*** | **-1.28** | **0.002** | **-2.08** | **-0.49** |
| | Similar | Anterior | 0.11 | 0.83 | -0.96 | 1.19 |
| | | Posterior | -0.24 | 0.72 | -1.54 | 1.06 |
| | Exception | Anterior | -0.13 | 0.75 | -0.99 | 0.72 |
| | | Posterior | 0.13 | 0.75 | -0.72 | 0.99 |
| Late | Anchor | Anterior | 0.54 | 0.27 | -0.43 | 1.50 |
| | | Posterior | -0.54 | 0.27 | -1.50 | 0.43 |
| | Similar | Anterior | 0.62 | 0.23 | -0.40 | 1.64 |
| | | Posterior | -0.62 | 0.23 | -1.64 | 0.40 |
| | Exception | Anterior | -0.22 | 0.47 | -0.82 | 0.39 |
| | | Posterior | 0.22 | 0.47 | -0.39 | 0.82 |

*Note.* * $p<0.05$
